# The coordination of innate and adaptive immunity in bacteria

**DOI:** 10.1101/2022.06.21.496935

**Authors:** Clemente F. Arias, Francisco J. Acosta, Federica Bertocchini, Miguel A. Herrero, Cristina Fernández-Arias

## Abstract

Bacteria have evolved a variety of innate and adaptive immune strategies to fight bacteriophage (phage) infections. Innate defenses (unspecific mechanisms directed against any phage infecting the cell) range from the identification and cleavage of the viral DNA by restriction nucleases to the suicidal death of infected host cells, an extreme solution that prevents the spread of the infection throughout the population. Adaptive immunity, on the other hand, involves the creation of an immune memory that targets specific phages in case of reinfection. It is obvious that not every infection leads to the suicide of the host cell or to the formation of immune memory against the infecting phage, so what determines the outcome of an anti-phage response? In this work, we suggest that the dynamic aspects of phage infections are key to addressing this question. We show that the rates of viral DNA replication and cleavage define functional categories of phages that differ in their susceptibility to the immune strategies evolved by bacteria. From this viewpoint, the combined action of diverse bacterial defenses would be necessary to reduce the chances of phage immune evasion. This perspective allows us to formulate simple molecular mechanisms that could account for the decision of infected cells to undergo suicidal cell death or to incorporate new phages into the immune memory. This work highlights the importance of dynamics to understand bacterial immunity and formulates explicit hypotheses that could inspire a new and original empirical approach to the study of phage/bacteria interactions.

## Introduction

The constant threat posed by bacteriophage (phage) infections has driven the development of a wide variety of immune mechanisms in bacteria [1–5]. Among the most common anti-phage defenses are the restriction-modification (RM) systems that detect and attack foreign DNA present in the bacterial cytoplasm [6, 7]. This immune strategy usually involves two types of enzymes: restriction nucleases that cleave DNA at specific sequences (known as restriction sites), and methyltransferases that modify the same sequences in the host DNA to avoid the “autoimmune” destruction of self DNA by nucleases [8, 9]. The breaks created by restriction nucleases in the viral DNA facilitate its subsequent digestion by other enzymes such as the RecBCD complex [10, 11]. The combined action of nucleases and methyltransferases ensures that only unmodified DNA is identified as non-self and destroyed by the cell.

Another defense strategy widespread in bacteria, known as abortive infection (Abi), consists of the suicidal death of infected cells before the completion of the phage replicative cycle, which prevents the spread of the infection to neighboring bacteria. [5, 12]. Although this strategy is encoded by an array of different molecular mechanisms, every Abi system requires two complementary functions: one that evaluates the evolution of the infection and one that kills the cell when the phage has escaped other bacterial defenses [12]. An example of this logic is the Rex system, one of the first Abi strategies to be described in the literature [13, 14]. This mechanism is activated by a sensor protein, RexA, capable of detecting protein-DNA complexes that appear in the bacterial cytoplasm during phage infections. RexA activates a second protein, RexB, that forms ion channels in the cell membrane, inhibiting bacterial growth and leading to the eventual death of the infected cell [15, 16].

Abi systems must be tightly controlled so that the suicidal death of the host cell only occurs after the phage has evaded other defense mechanisms but before it has had time to complete its lytic cycle. Achieving such precise timing is far from trivial. It is usually assumed that RM systems must operate at the early stages of the infection and Abi systems in later stages [12, 17, 18]. However, the mechanisms allowing bacteria to unambiguously discriminate early from late phases of phage infections remain largely unexplained.

The previous immune mechanisms must also be coordinated with other anti-phage defenses. Both RM and Abi systems can be defined as “innate” mechanisms: they do not keep a memory of past infections and are unspecific, i.e. they can target any phage infecting the cell. Remarkably, bacteria also resort to “adaptive” immunity: the CRISPR system (present in about 40% of all types of bacteria [19]) creates immune memory against previous phage infections. The number and diversity of known CRISPR systems have steadily grown since the seminal works that led to its discovery [20–24]. In spite of their diversity, they all respond to a similar underlying logic. A typical CRISPR immune response consists of three main steps: adaptation, expression, and interference. At the adaptation stage, CRISPR-associated proteins (Cas) bind to the DNA of an infecting phage and cleave a small fragment that is subsequently inserted into the CRISPR array, becoming a new spacer. The expression phase occurs during reinfections and consists of the transcription of the CRISPR array into precursor CRISPR RNA, which then undergoes enzymatic cleavage to yield mature CRISPR RNAs that usually contain a single spacer sequence [25]. At the interference stage, these RNAs bind their target nucleotide sequences in the phage genome and recruit Cas nucleases that cleave the phage DNA, thus preventing its replication [26].

The length of the CRISPR array in bacterial genomes ranges from a few dozen to a few hundred spacers [27]. Intuitively, it seems that increasing the number and variety of spacers should make bacteria less vulnerable to phages. However, the number of spacers is also subject to an opposite selective force. As we noted above, the CRISPR array is usually transcribed as a single unit and then cleaved into individual RNA molecules corresponding to single spacers [24]. This means that all the spacers present in the CRISPR memory are recruited in case of infection, even those that cannot target the phage infecting the cell at that moment. An excess of spacers would therefore dilute the number of functional CRISPR units against the ongoing reinfection, reducing the overall effectiveness of the system [28]. Furthermore, the predominance of deletions over insertions in bacterial genomes tends to reduce their size [29], which imposes an additional pressure against larger CRISPR arrays.

Given the ubiquity of phages, the fact that some spacers remain in bacterial populations for very long periods of time suggests that not all episodes of infection result in the inclusion of new spacers into the CRISPR library [26, 30]. Otherwise, owing to the size constraints discussed above, recent infections would displace earlier ones [31, 32], hindering the maintenance of long-term memory. This raises the question of how bacteria decide whether or not a given phage or plasmid DNA has to be integrated into their CRISPR arrays. To the best of our knowledge, this issue remains largely unsolved.

This work is motivated by the following questions: How do bacteria coordinate the RM, Abi, and CRISPR systems in the course of anti-phage responses? Are these systems redundant or do they target different phage strategies? In what circumstances is a new phage incorporated into the CRISPR array?

To address the previous questions, we formulate simple mathematical models of bacterial immune responses. These models suggest that the relative rates of phage DNA replication and cleavage suffice to characterize the outcome of phage infections, defining functional categories of phages that differ in their susceptibility to bacterial immune mechanisms. Within this framework, RM, Abi, and CRISPR systems play complementary roles in bacterial immunity, targeting different phage strategies. We show that the coordination between these anti-phage mechanisms during infections naturally emerges from the dynamics of phage/bacteria interactions. Finally, we suggest that those dynamics give rise to a simple molecular mechanism that would allow bacteria to decide if a new phage must be included in their CRISPR library.

## Results

### The role of restriction nucleases during phage infections

Phages may use two alternative strategies to infect bacterial populations. Lysogenic phages integrate their genome into the bacterial chromosome and replicate without lysing the host. In contrast, lytic phages destroy the host cell to release hundreds of new phages that can infect nearby bacteria [33, 34]. Lytic infections, the focus of this work, are usually described by means of qualitative models that represent the key discrete events of the infection (Fig. 1). Although these models are useful to describe the interactions between phages and their hosts, they are insufficient to fully understand the progression of phage infections. The fate of the host cell depends on the balance between the effectiveness of its immune mechanisms and the ability of the phage to subvert those mechanisms. However, from the conceptual model shown in Fig. 1 it is impossible to deduce in what circumstances lytic phages prevail over the host’s immune defenses or how infected cells decide when they must resort to suicide.

**Figure 1.**
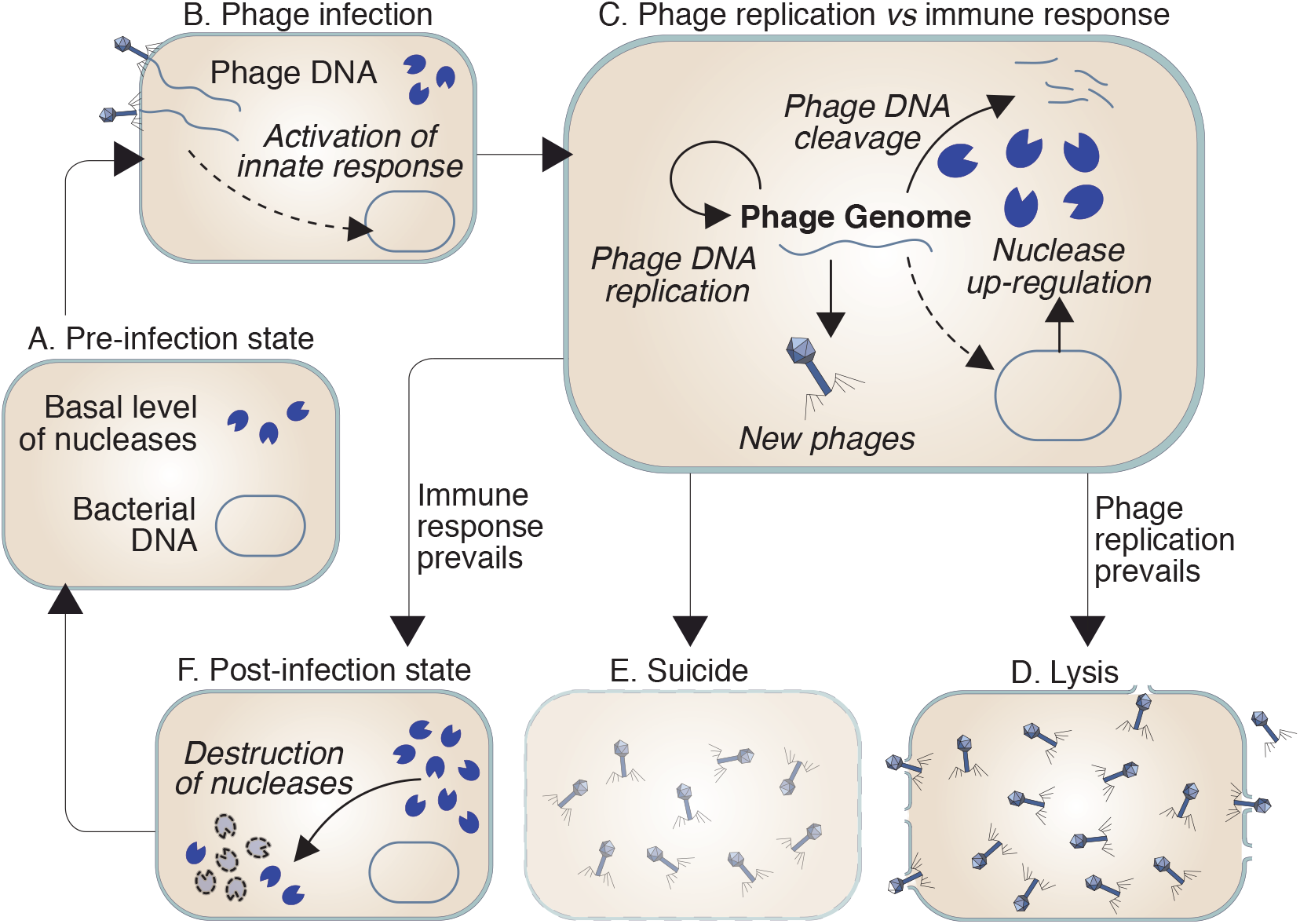
Standard representation of the role of nucleases in lytic infections [1]. A) In the absence of infection, bacteria are free from viral DNA and may display basal levels of restriction nucleases capable of cleaving non-self DNA. B) In case of infection, the host detects the presence of phage DNA and triggers an “innate” immune response that includes the upregulation of restriction nucleases. C) The balance between the replication of the viral DNA and its destruction by bacterial nucleases determines the eventual outcome of the infection. D) If the phage evades the host’s immune response, the bacterial cell dies and releases the newly formed phages to the extracellular space. E) Alternatively, infected cells may undergo suicidal cell death to prevent the spread of the infection. F) If the host cell survives, it must deactivate its anti-phage defenses and return to the pre-infection state, which involves the downregulation of the nucleases produced during the infection and also the clearance of the remnants of viral DNA from the cytoplasm.

To answer these questions, the qualitative description of nuclease responses has to be translated into quantitative terms. To do that, we will begin by modeling the interaction between nucleases and the viral DNA during phage infections. We will use the number of phage genomes in the bacterial cytoplasm as an indicator of the progression of the infection. We will assume that if this number reaches a critical value, then the host cell dies; on the contrary, the host cell effectively controls the infection if this number falls below a certain minimum. To model the dynamics of the phage, we will further assume that phage genomes replicate exponentially and disappear by the enzymatic action of nucleases according to the following equations:

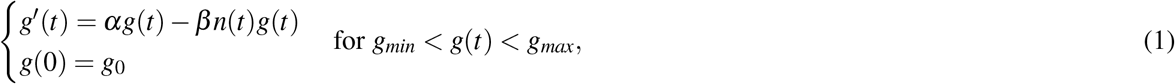

where *g*(*t*) and *n*(*t*) are the number of phage genomes and of nucleases at time *t* respectively, *g*_0_ is the initial number of phage genomes that infect the host cell, *g*_*min*_ is the lower threshold that determines phage viability, and *g*_*max*_ is the number of phage genomes that causes the lysis of the host cell. Positive parameters *α* and *β* represent the phage DNA replication and cleavage rates respectively.

With regard to nucleases, we remark that their behavior during anti-phage responses is analogous to the clonal expansion and contraction of T cells during adaptive immune responses in vertebrates. The number of nucleases in the bacterial cytoplasm “expands” when the cell detects the infection and “contracts” once the infection is resolved (Fig. 1). This analogy is not merely superficial but responds to an identical functional strategy. Both nucleases and effector T cells must remain inactive under normal conditions. Their numbers must rapidly increase (through activation and cell division in the case of T cells and through upregulation in the case of nucleases) to fight an ongoing infection. After the threat is neutralized, the defenses are de-activated (through apoptosis in T cells and downregulation of nucleases in bacteria [35]), which restores the pre-infection state. T cells and nucleases can be described as “elastic” systems since they change in size in response to an external stimulus (the infection) and return to their original value when that stimulus disappears. In previous works, we have shown that this feature of T cells can be naturally modeled by means of second-order differential equations (see [36] and [37] for further details on this point). Exploiting the analogy of nucleases with T cells, we will use this approach to model the interaction between nucleases and phage DNA as follows:

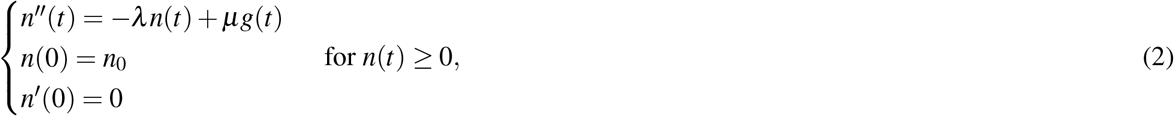

where *g*(*t*) and *n*(*t*) are the number of phage genomes and nucleases at time *t* respectively, *n*_0_ is the number of nucleases in the bacterial cytoplasm before the infection, and *k, λ, α* and *β* are positive parameters. We assume that the number of nucleases in the absence of infection is at a homeostatic equilibrium (hence the condition *n*^*l*^(0) = 0).

Putting equations 1 and 2 together, the nuclease response to a phage infection can be modeled by the following system of differential equations:

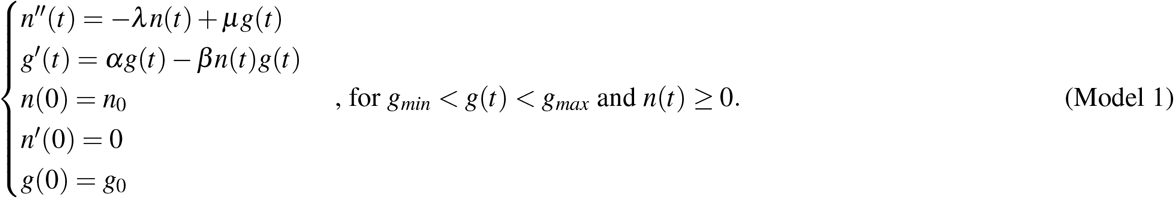

As should be expected, Model 1 captures the elastic nature of nucleases: they proliferate in response to the presence of the phage and disappear after the infection is neutralized (see 1 in Fig. 2.A). Importantly, this model also reveals key aspects of phage infections that are not evident in its qualitative counterpart (as outlined in Fig. 1). In particular, it suggests that phages may adopt two alternative strategies to evade nuclease responses. Phages with very high DNA replication rates prevail by lysing the host cell before the nucleases expand sufficiently to fight the infection (see 2 in Fig. 2.A). Less intuitively, phages with very low DNA replication rates give rise to “chronic” infections characterized by successive cycles of DNA replication and degradation (see 3 in Fig. 2.A). In this case, the progressive accumulation of viral particles in the bacterial cytoplasm would affect the fitness of the infected cell and could even lead to its eventual death.

**Figure 2.**
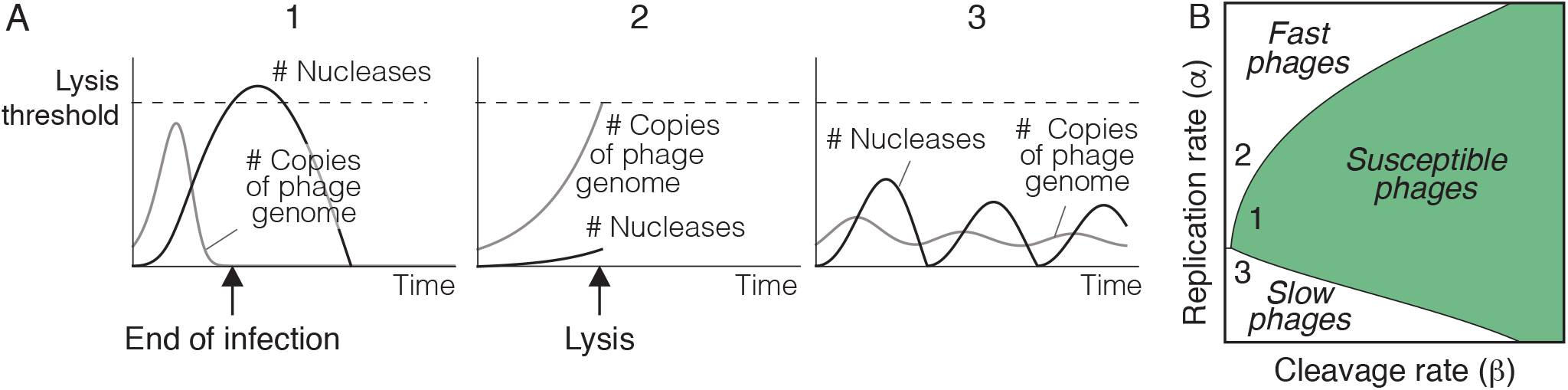
Possible outcomes of the interaction between viral DNA and restriction nucleases. A) Model 1 gives rise to three types of outcomes: 1) phages can be successfully eliminated by the expansion of nucleases (condition *g*(*t*) *< g*_*min*_), 2) phages can kill the cell before nucleases have had time to expand (condition *g*(*t*) *> g*_*max*_), and 3) nucleases and viral DNA can oscillate and give raise to a sustained infection that eventually kills the host cell (when none of the previous conditions is fulfilled). B) The outcomes of Model 1 depend on the parameters used in the numerical simulations. This model defines an infection space, characterized by the rates of viral DNA replication and cleavage (parameters *α* and *β* in Model 1 respectively). Within this space, phages can be classified into three functional categories: 1) Phages susceptible to the action of nucleases as the one shown in A1, 2) phages with high replication rates that outrun nucleases (like in A2), and 3) phages with low replication and cleavage rates that induce oscillations in the number of nucleases (like in A3). Based on the dynamics shown in A, we will label these types of phages as susceptible, fast, and slow respectively. The details of the simulations are provided in the Methods section.

Model 1 defines an “infection space” in which the outcome of anti-phage responses is fully characterized by the rates of phage DNA replication and degradation by restriction nucleases (parameters *α* and *β* respectively). Within this space, phages can be classified into three functional categories that we will label as susceptible (those that can be eliminated by restriction nucleases), fast (those that outrun the expansion of nucleases), and slow (those that cause chronic infections) (see Fig. 2.B). We remark that this classification naturally emerges from a very simple formalization of the standard description of phage infections shown in Fig. 1.

### Bacterial suicide: a strategy against fast phages

In this section, we will model the dynamics of typical Abi systems to show that they confer bacteria protection against fast phages. As discussed above, Abi systems must sense the progression of the infection and kill the cell if the phage cannot be neutralized by other immune defenses [12]. At the same time, Abi proteins must be tightly suppressed under normal conditions to prevent the death of uninfected bacteria [12]. For this reason, the effector mechanisms of Abi systems are normally dormant proteins that only operate when a phage infection is detected in the bacterial cytoplasm [17]. The sensor mechanisms of Abi systems recognize a wide variety of stimuli, ranging from the formation of intermediates of phage replication to the disruption in the expression of host genes caused by the activity of the phage [12]. We will use this last alternative as a model to simulate the behavior of an Abi with the following features: i) It monitors the levels of a host protein *s* whose expression decreases during the infection; ii) It triggers the death of the cell if this protein disappears from the bacterial cytoplasm. The dynamics of this protein can be simply modeled as follows:

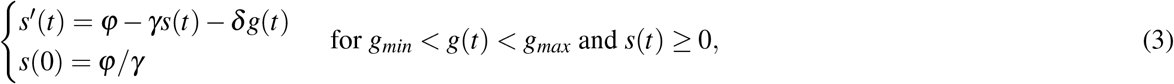

where *ϕ, γ*, and *δ* are positive parameters. This model simulates the dynamics of a protein that remains at homeostatic equilibrium under normal conditions (*s* = *ϕ/γ*) and disappears as a consequence of the phage activity in case of infection. Including equation 3 in Model 1, the simultaneous action of RM and Abi systems can be described by the following system of differential equations:

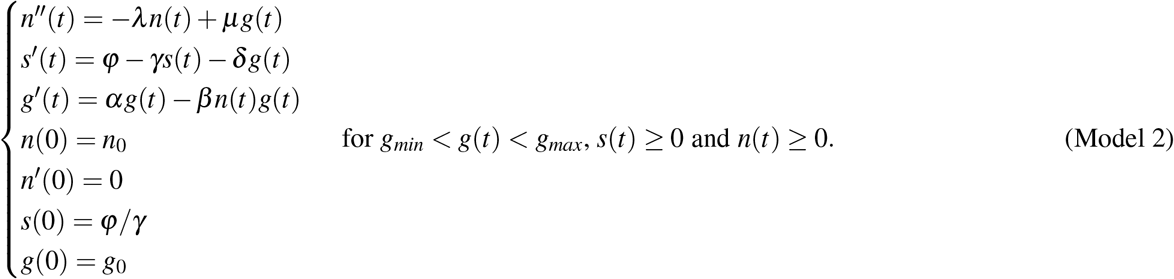

The three possible results of phage infections shown in Fig. 1 can be explained as alternative outcomes of this model. The phage is neutralized if *g*(*t*) ≤ *g*_*min*_, the host cell is killed by the Abi effector molecules if *s*(*t*) ≤0, and the phage survives in the rest of the scenarios. The outcome of each anti-phage response is determined by the first of those conditions to be fulfilled in the course of the infection. According to the behavior predicted by our model, bacterial suicide predominantly occurs in response to infections by fast phages (Fig. 3.A), which implies that Abi systems can kill the host cell before these phages can complete their lytic cycles. It is remarkable that, on the other hand, restriction nucleases neutralize slower phages before the Abi systems have time to induce the death of the host (Fig. 3.A). In these cases, the disappearance of the phage prevents Abi sensors from activating the effector mechanisms that kill the host cell.

**Figure 3.**
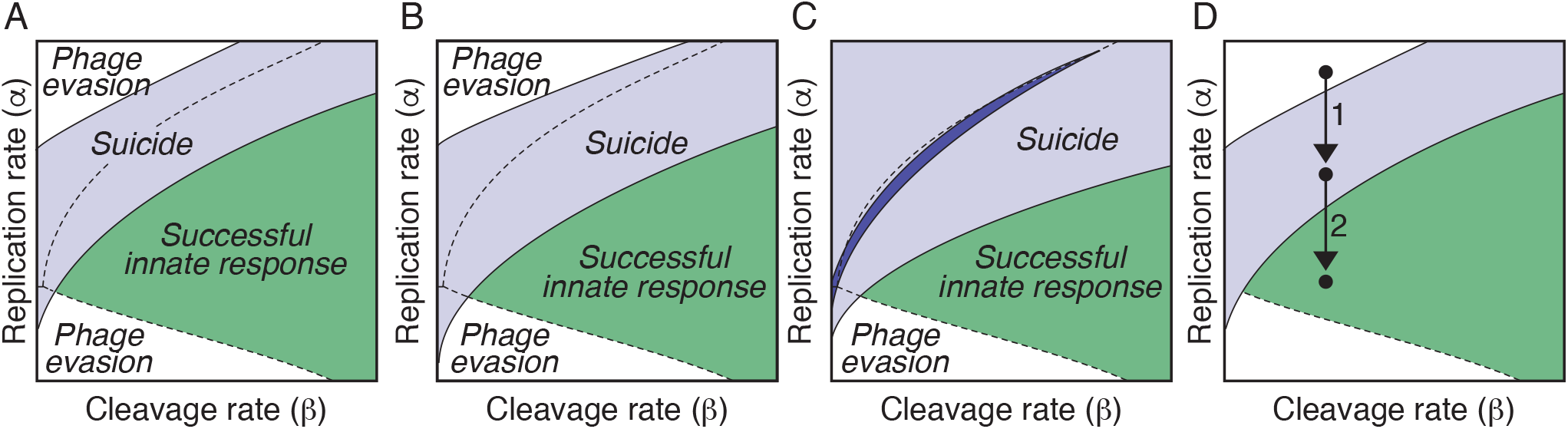
Coordination of innate immune responses in bacteria. A) Numerical simulations of Model 2 show that bacterial suicide predominates in a region of the infection space that overlaps with fast phages. This implies that Abi systems can neutralize phages that are not susceptible to restriction nucleases. B) The suicide region produced by an alternative Abi system (described by equations 4) is similar to that shown in A. C) Changing the values of the parameters of equation 3 in Model 2 gives raise to different suicide regions (shown in clear and dark blue respectively). D) The state of dormancy induced by Abi systems in infected cells reduces the rate of phage replication. As a consequence, host cells can resort to suicidal death against phages that would otherwise escape the action of bacterial immunity (1). A sufficiently large reduction in the rate of viral DNA replication by Abi systems may allow for nucleases to eliminate fast phages, preventing both the spread of the infection and the suicide of the host cell (2). The details of the simulations are provided in the Methods section.

The previous results hint at a new explanation for the role of bacterial suicide during phage infections. Abi systems would not target the same phages as RM systems, and would not be activated when other immune mechanisms fail to control the infection. Quite the opposite, Abi and RM systems would operate simultaneously but they would be programmed to target different categories of phages. RM would neutralize susceptible phages that are beyond the control of Abi systems, whereas host cells would undergo suicidal cell death in infections by fast phages that restriction nucleases cannot neutralize.

In terms of Model 2, RM and Abi systems occupy different regions of the infection space (Fig. 3.A). We remark that the shape of the Abi region is determined by the dynamics of its sensor mechanism. To clarify this point, let us consider a different Abi system whose sensor mechanism detects the presence of phage DNA. Such a mechanism can be described by the following equations:

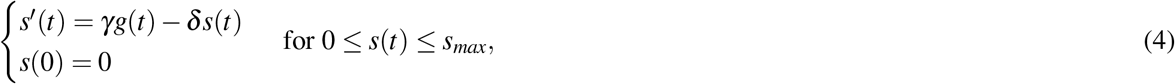

where *γ* and *δ* are positive parameters, *s*(*t*) is the sensor protein, and *g*(*t*) is the number of phage genomes in the bacterial cytoplasm. According to this model, the concentration of the sensor protein increases proportionally to the number of genomes in the bacterial cytoplasm and decreases at a rate *δ*. We assume that this protein triggers the death of the host cell whenever its concentration rises above a threshold value *s*_*max*_. Substituting equations 3 by equations 4 in Model 2 yields similar results (see Fig. 3.B), which shows that Abi systems based on the detection of phage DNA can also target fast phages.

Regardless of their particular mechanistic details, the range of action of Abi systems must be subject to a tradeoff between two alternative constraints. On the one hand, these systems should not kill the host cell if RM systems suffice to control the infection and on the other, they should accelerate the suicidal death of cells infected by fast phages. Within the framework of Model 2, these constraints imply that the region occupied by Abi systems in the infection space should minimize the intersection with susceptible phages while maximizing the one with fast phages. This could be achieved by varying the kinetic parameters of the sensor enzymes used by Abi systems (Figs. 3.C). These parameters could be modulated by natural selection to fine-tune the configuration of Abi systems (i.e. the phages that should trigger the suicide of infected cells) depending on the ecology of each bacterial species.

Abi systems have also been reported to induce a state of dormancy in infected cells by reducing their metabolic rate [5, 12, 17]. This behavior is usually interpreted as an alternative to suicidal death intended to “buy time” for the action of other immune mechanisms [12]. Our model suggests a different explanation for this phenomenon. Reducing the metabolic activity of infected cells would lower the rate of phage replication, forcing fast phages into the Abi region (Fig. 3.D). This would eliminate phages that would otherwise kill the host cell and infect neighboring bacteria (Fig. 3.D). Lowering the phage replication rate could also make fast phages susceptible to the action of RM systems, preventing the suicidal death of the cell (Fig. 3.D). From this viewpoint, dormancy and suicide would be alternative results of the same mechanism operating against phages with different replication and destruction rates. Incidentally, this approach would also account for the anti-phage effect of other mechanisms that drastically inhibit phage replication by slowing down the activity of infected cells [38].

### The CRISPR system: a bacterial strategy against slow phages

In the previous section, we have seen that RM and Abi systems would fail to protect bacteria against slow phages (Fig. 3). We suggest that CRISPR systems would be an adaptation to fight this category of phages. In agreement with this view, we will next show that adaptive immune responses could eliminate slow phages. In the next section, we will present a hypothetical molecular mechanism that would allow bacteria to discriminate slow phages and incorporate them into the CRISPR array.

During the interference stage of CRISPR responses, Cas proteins cleave the phage DNA at specific sites recognized by CRISPR RNAs. As occurs with restriction nucleases, Cas nucleases must expand after the detection of a pathogen, and contract once the phage is eliminated. Therefore, their dynamics can be described by the following equations:

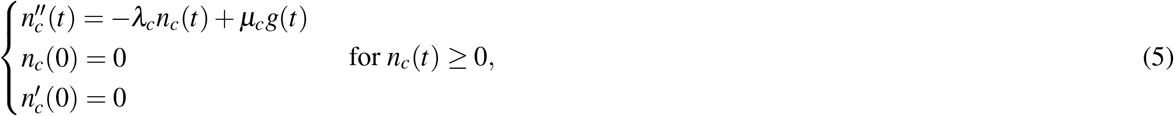

where *λ*_*c*_ and *µ*_*c*_ are positive parameters. We assume that Cas nucleases are only expressed in case of infection (hence, the initial conditions *n*_*c*_(0) = 0 and *n*^*l*^_*c*_(0) = 0).

During infections by phages present in the CRISPR library, Cas and restriction nucleases cooperate to cleave the viral DNA. The combined action of restriction and CRISPR nucleases on the phage DNA can be modeled by including equations 5 in Model 1:

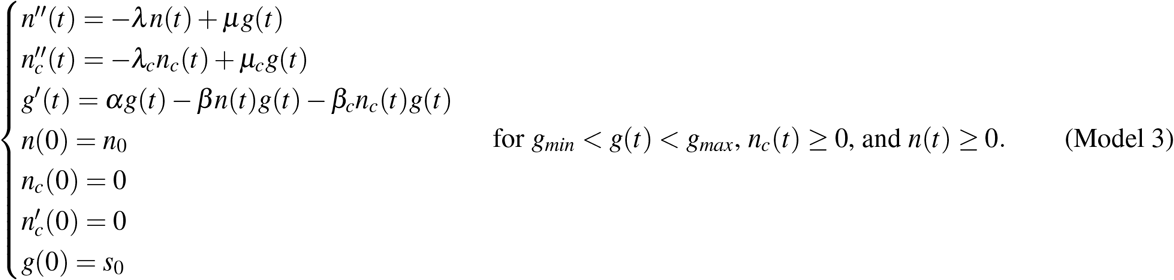

Numerical simulations of this model for slow phages show that although the CRISPR system would not always prevent the death of the host cell during infections by slow phages, it could neutralize some infections that would otherwise escape the control of nucleases (Figs. 4.A). The strategy of using the CRISPR systems against slow phages would thus reduce the regions of the infection space available for viral immune evasion (Figs. 4.B-C). The combined action of RM, Abi, and CRISPR systems would force the selection of phages in two different directions. First, phages would be selected toward lower nuclease-mediated cleavage rates. This selective force drives the frequent mutations that modify the restriction sites in the viral DNA, reducing its likelihood of nuclease-mediated recognition and cleavage [39] (Fig. 4.D). According to our model, this strategy alone would not ensure that phages can successfully evade bacterial immunity. Abi and CRISPR systems would impose a second selective pressure on the rates of DNA replication. Depending on their DNA cleavage rates, phages would be forced to replicate either faster or slower to avoid immune destruction (Fig. 4.D).

**Figure 4.**
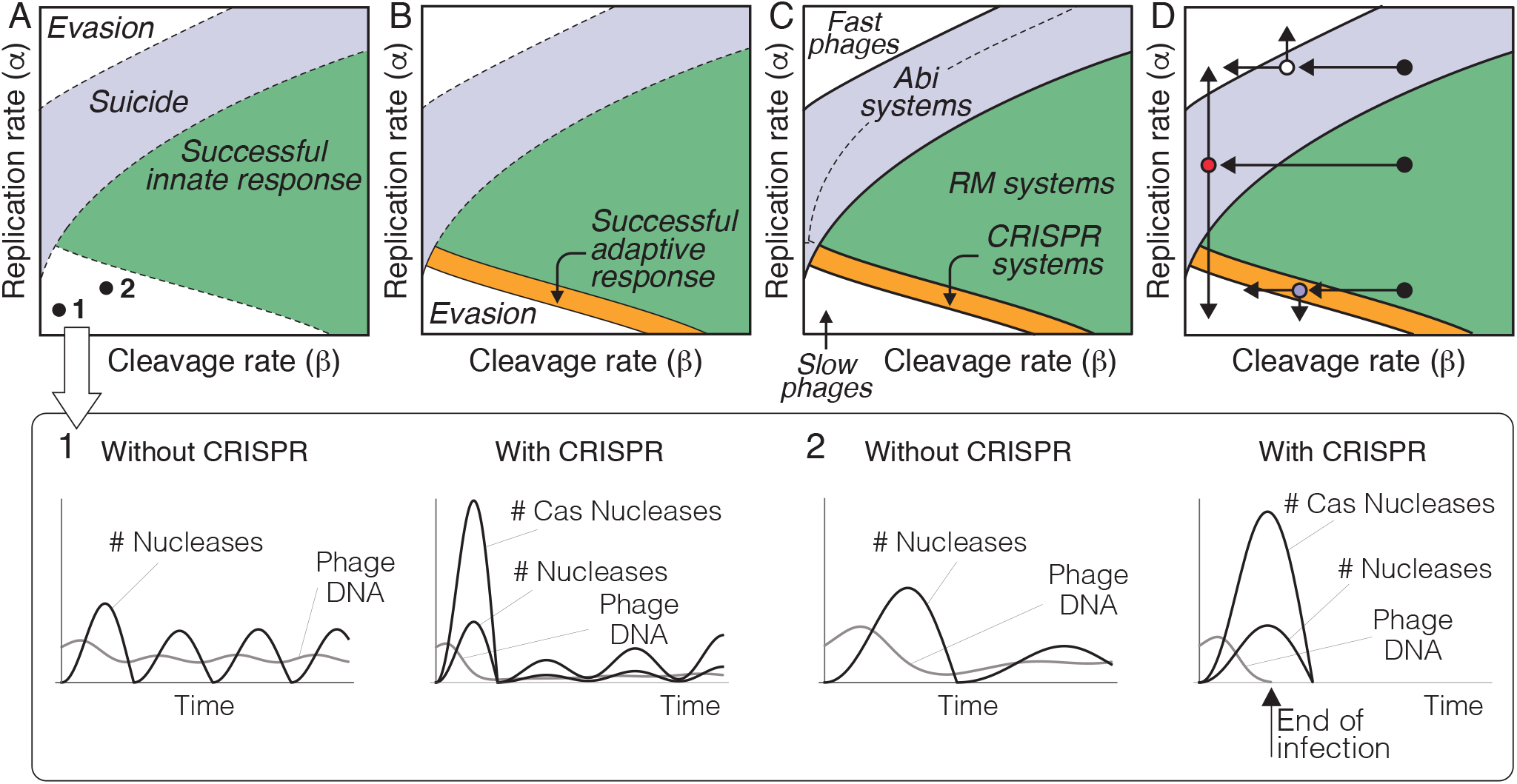
Role of the CRISPR systems in anti-phage responses. A) Effect of CRISPR spacers on slow phages. The activation of CRISPR nucleases increases the rate of phage DNA degradation. This does not affect some phages that can still progress with the infection and cause the eventual death of the host cell (1). In other cases, the increase in the rate of DNA cleavage due to CRISPR nucleases can entail the end of the infection (2). B) CRISPR spacers create a new region in the infection space where the host cell can neutralize slow phages (shown in orange). Slow phages that can still evade the action of CRISPR spacers are resistant to bacterial adaptive immunity. C) Regions occupied by innate and adaptive immune mechanisms in the infection space. D) Selective pressures imposed on phages by bacterial immune mechanisms. Reducing the rate of DNA cleavage may not suffice for phages to evade bacterial immunity (black circles). Although they may be less vulnerable to restriction nucleases, they can become the target of Abi (white circle) or CRISPR systems (blue circle). Fast phages are forced to further reduce their rate of DNA cleavage or to replicate faster (white circle), whereas slow phages must reduce both the rates of DNA replication and destruction to escape bacterial immunity (blue circle). The only alternative for susceptible phages is to greatly increase or decrease their replication rates (red circle). The details of the simulations are provided in the Methods section.

### How do bacteria decide to include new phages in the CRISPR array?

In the previous section, we have shown that the CRISPR systems would eliminate some slow phages capable of evading bacterial innate immunity. To understand how could CRISPR systems exclusively target slow phages, we begin by remarking that the creation of new CRISPR spacers requires the formation of stable complexes between Cas proteins Cas1 and Cas2 during the adaptation stage of the CRISPR immune response [25]. Cas1-Cas2 complexes then interact with the fragments of phage DNA that will be added to the CRISPR library [40]. This interaction obviously requires that fragments of phage DNA are still present in the host cytoplasm after the formation of Cas1-Cas2 complexes. Based on this observation, we hypothesize that the incorporation of new spacers into the CRISPR array is determined by the permanence of phage DNA in the bacterial cytoplasm. This mechanism relies on two basic assumptions. The first one is that the formation of Cas1-Cas2 complexes is delayed with respect to the initiation of the innate anti-phage responses, which implies that when these complexes appear in the cytoplasm the phage DNA is already undergoing nuclease-mediated degradation. Our second assumption is that Cas1-Cas2 complexes can only create new spacers if they encounter suitable fragments of phage DNA in the host cytoplasm (see Fig. 5). Both assumptions are supported by the observation that Cas1-Cas2 complexes create new spacers using the debris that appears from the degradation of the viral DNA by nucleases [25].

**Figure 5.**
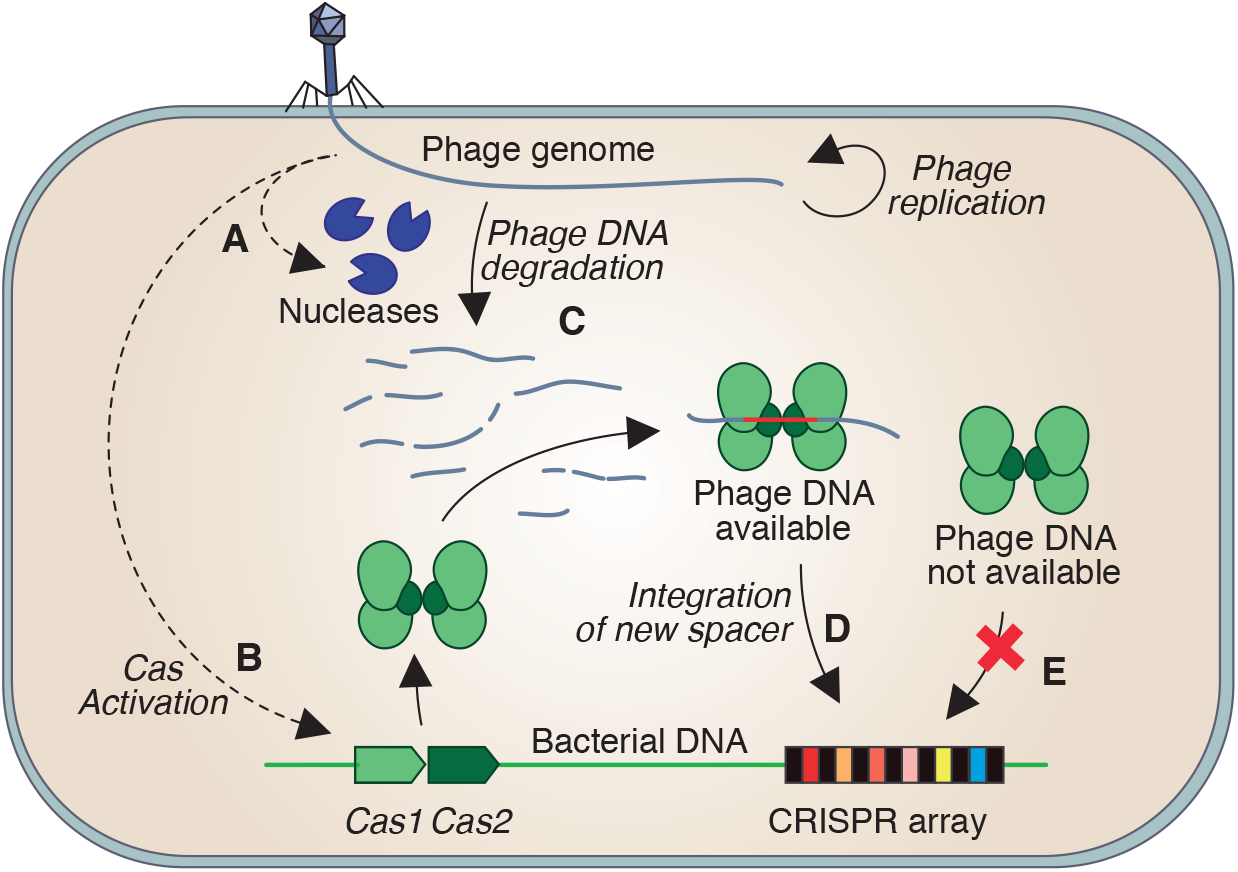
Hypothesized model of the inclusion of new phages in the CRISPR memory. The detection of a phage infection up-regulates restriction nucleases that cleave the viral DNA at specific restriction sites, fostering its subsequent digestion by other bacterial enzymes (A). Our model assumes that Cas enzymes are activated in the late stages of the innate response to the infection (B), once the viral DNA has already undergone a certain degree of nuclease-mediated degradation (C). New spacers against the infecting phage can only be created if fragments of its DNA are still present in the cytoplasm after the formation of Cas1-Cas2 complexes (D). Otherwise, Cas nucleases have no substrate to act, which prevents the incorporation of the phage into the CRISPR array (E).

Now, according to Model 2, the time needed by nucleases to neutralize susceptible phages is not homogeneous. Their action is faster against phages with greater rates of DNA cleavage and replication (Fig. 6.A). In contrast, the permanence of DNA from slow phages in the bacterial cytoplasm is much longer (see Figs. 6.A and 2.A). In consequence, the second assumption of the model outlined in Fig. 5 would only be verified by slow phages. Therefore, the delay of Cas activation relative to the velocity of action of restriction nucleases would suffice to discriminate between slow and fast phages and to exclude the latter from the CRISPR library (Fig. 6.B).

**Figure 6.**
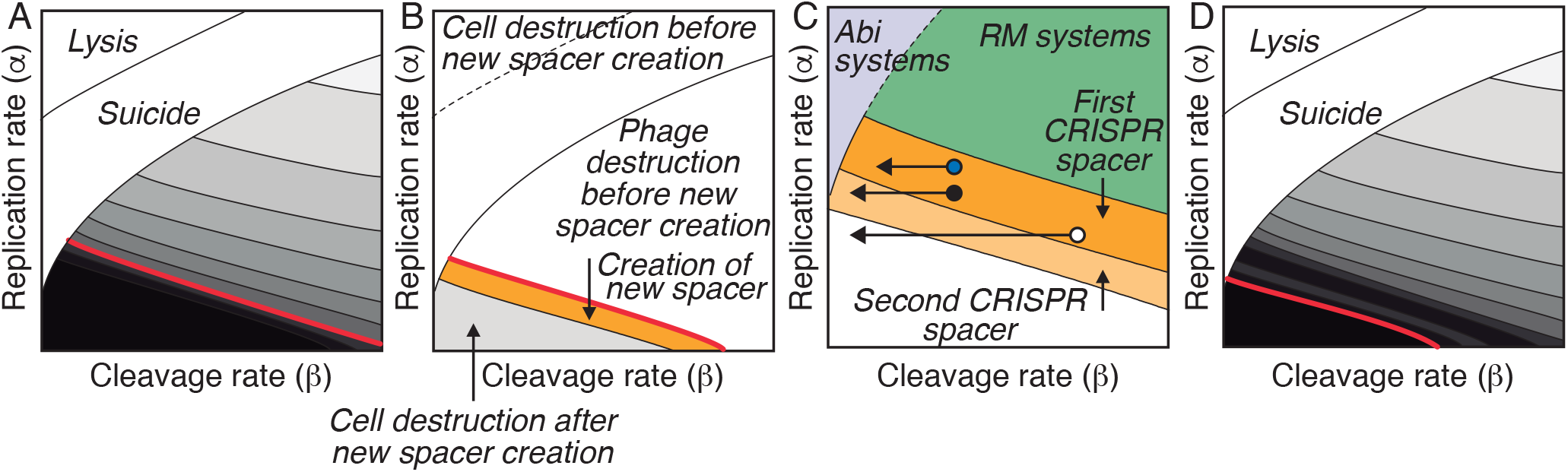
Functional consequences of CRISPR. A) Contour plot of the permanence of phage DNA in the cytoplasm of infected bacteria. Different shades of gray indicate the time needed for nucleases to remove the phage DNA from the bacterial cytoplasm (lighter colors indicate a faster elimination of phage DNA.) Higher rates of DNA replication and cleavage reduce the time needed by nucleases to eliminate the phage DNA. The black region corresponds to slow phages that evade the action of nucleases. We hypothesize that the time it takes for nucleases to remove phage DNA from the cytoplasm of infected bacteria provides a mechanistic criterion to include new phages in the CRISPR library. The late activation of the Cas enzymes (indicated by the red line) would constrain the creation of new spacers against susceptible phages as shown in B. B) Some susceptible phages would be eliminated before the formation of Ca1-Cas2 complexes (indicated by the red line). Fast phages, on the other hand, would kill the cell before the activation of the Cas enzymes. During infections by slow phages, Cas1-Cas2 would have enough time to create new spacers and therefore include the infecting phage in the CRISPR array (orange region). Although host cells could also incorporate very slow phages into the CRISPR library (gray region), they would be eventually killed by the infection, preventing the future use of those CRISPR spacers (death host cells cannot be reinfected). C) Reinfections by phages that have changed their rates of DNA cleavage have three possible outcomes: the host cell can eliminate the phage (blue circle), it can create a new spacer against the same phage (black circle), it can be killed by the phage (white circle). D) In case of reinfections by phages that have not changed their rate of DNA cleavage, the action of CRISPR nucleases accelerates the elimination of the infection, so the permanence of the phage DNA is shorter than during previous infections. The Cas enzymes would not have enough time to create new spacers against these phages. (Lighter shades of gray indicate shorter permanence time of the phage DNA in the bacterial cytoplasm.) The details of the simulations are provided in the Methods section.

This model has an interesting functional implication related to the behavior of the CRISPR systems during reinfections. Should bacteria add new spacers against phages that are already represented in their CRISPR array in case of reinfection? This issue concerns an important aspect of the bacterial immune memory that critically affects the function of CRISPR systems. Apparently, creating new spacers against the same phage during each episode of reinfection would reduce the utility of the system by decreasing the diversity of spacers in the CRISPR library. This would be especially so in the case of phages that cause frequent reinfections, which would soon displace other phages and dominate the bacterial immune memory [41]. We will next show that the mechanism of immune formation hypothesized above provides a straightforward solution to this issue.

To that end, we will consider two different scenarios of reinfection. First, let us suppose that reinfecting phages present in the CRISPR library are less susceptible to the host’s nucleases than in previous infections (owing, for instance, to mutations in the restriction sites). This would imply a lower rate of viral DNA degradation by nucleases. In this case, the additional cleavage of the phage DNA by the CRISPR enzymes could contribute to neutralizing the infection (Fig. 6.C). Alternatively, the combined action of restriction and CRISPR nucleases could be insufficient to eliminate the phage, which would kill the host cell (Fig. 6.C). A third possibility is the creation of new spacers against the phage. The presence of several spacers against the phage would further increase the rate of destruction of its DNA, which could shift the balance toward the host cell and terminate the infection (Fig. 6.C). Therefore, the creation of new spacers against phages already represented in the CRISPR array could be advantageous for bacteria under some circumstances. In the case of reinfection by similar phages, only those cases where the phage susceptibility to nucleases has changed would result in adding new a spacer in the CRISPR library.

Let us now consider a second scenario in which the rate of DNA cleavage has not changed since previous contacts between the phage and the host cell. In this scenario, the already acquired CRISPR memory will add to the innate immune response to facilitate the elimination of reinfecting phages. The rapid disappearance of the phage DNA from the bacterial cytoplasm would prevent the creation of new spacers during this type of reinfection (Fig. 6.D).

In summary, our model of immune memory formation in bacteria accounts for the decision to incorporate a phage into the CRISPR array the first time it infects a host cell. The same model provides a functional criterion to include new spacers against phages already present in the CRISPR library during reinfections. Finally, it explains how could CRISPR systems target slow phages, which are precisely the type of infections that cannot be neutralized by innate immune mechanisms.

## Discussion

In this work, we use a simple mathematical description of the standard models of phage/bacteria to gain insight into the dynamics of phage infections. This approach reveals unexpected features of anti-phage defenses that could hardly be understood from their qualitative descriptions. It suggests new and testable hypotheses about the coordination of bacterial immune systems: Fast and slow phages, resistant to restriction nucleases, would be the targets of Abi, and CRISPR systems respectively. Accordingly, the inclusion of new spacers in the CRISPR array would be biased towards slow phages.

Mathematical models cannot prove or disprove the validity of these hypotheses, a task that requires an empirical approach. What they prove, however, is that the dynamics of anti-phage systems, usually neglected in the literature, could account for key aspects of bacterial immunity. The importance of dynamics is explicitly acknowledged in the explanation of other bacterial mechanisms. For instance, it is widely accepted that the function of toxin-antitoxin systems depends on the different rates of degradation of two molecules, a toxin that can kill the host cell and an antitoxin that protects the host from this effect [42]. If protein synthesis is halted (owing to nutritional or environmental stress [43]), the antitoxin disappears from the bacterial cytoplasm before the toxin, which has a lower degradation rate [44]. In these circumstances, the toxin can no longer be neutralized and the host cell dies. The differences in the rates of degradation of the toxin and the antitoxin are obviously crucial to understanding the logic of this mechanism.

Our results suggest that taking into account the rates of activity of the molecules involved in anti-phage defenses could also be necessary to understand the coordination of anti-phage defenses. This approach sets the ground for a new framework for bacterial immunity in which phages and bacteria can be viewed as competing to occupy an infection space. Within this space, characterized by the rates of phage replication and destruction, phages would exploit the regions that are beyond the reach of bacterial defenses. In their turn, RM, Abi, and CRISPR systems would be designed to minimize the extent of those regions. We do not intend to provide an exhaustive and accurate description of the infection space, whose configuration depends on the molecular details of the immune mechanisms implemented in individual bacteria. Instead, we use very simple descriptions of anti-phage mechanisms to illustrate the explicative power of the dynamic aspects of anti-phage mechanisms. This approach provides an original perspective to interpret the interactions between phages and bacteria and paves the way to a better understanding of bacterial immunity.

## Acknowledgments

F.B. and C.F.A. are grateful to the Roechling Foundation for its support. Cr.F.A. was partially supported by the FCT grant no. EXPL/BIA-BIO-0644/2021.

## Methods

The numerical simulations of the models were performed with Wolfram Mathematica ^@^. To elaborate the figures of the infection space, the models were run for a discrete set of values of parameters *α* and *β*. For each set of parameters, the outcome of the simulations of Model 1 was determined by the following conditions: 1) if *g*(*t*) ≤ *g*_*min*_, the infection is controlled, 2) if *g*(*t*) ≥ *g*_*max*_, the host cell is lysed, and 3) otherwise the infection is chronic, which may entail the eventual death of the host cell. The outcome of the simulations of Model 2 was subject to an additional condition, imposed by the Abi sensors: if *s*(*t*) ≥ *s*_*max*_, the infected cell undergoes suicidal death. Based on the results of each simulation, the infection space was qualitatively described as shown in Fig.7.

**Figure 7.**
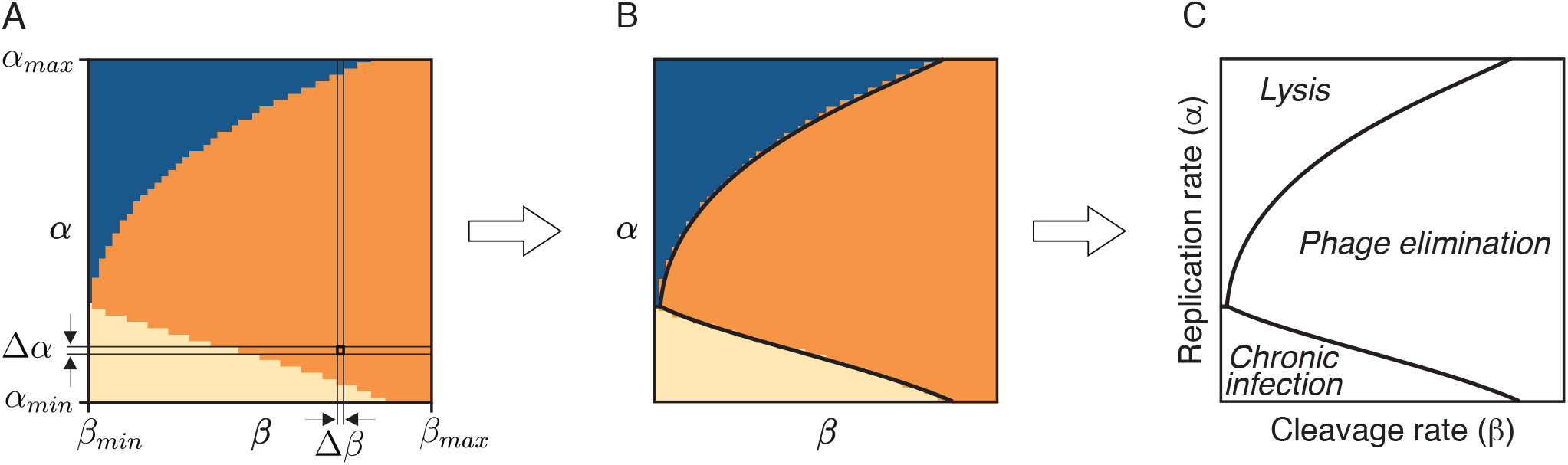
Characterization of the infection space from the simulations of Model 1. A) Numerical simulations of refeq1 have three possible outcomes: cell lysis (blue), control of the infection (orange), and chronic infection (light yellow). The simulations were performed for a range of values of parameters *α* and *β*, from a minimum (*α*_*min*_ and *β*_*min*_ respectively) to a maximum value (*α*_*max*_ and *β*_*max*_ respectively) with steps Δ*α* and Δ*β* respectively. B)These simulations give rise to a discrete version of the infection space. C) For the sake of simplicity, the figures show a representation of the boundaries between the different regions of the infection space. The rest of the figures in the text were constructed with the same logic.

In this work, we do not intend an exhaustive exploration of the parameter space or the sensitivity analysis of the models that describe anti-phage responses in bacteria. The values of the parameters used in the numerical simulations of these models were chosen to illustrate the behaviors that emerge from the dynamics of bacterial immune mechanisms. The specific values shown in the figures were the following:

- Figure 2. Numerical simulations of Model 1 for *λ* = 1, *µ* = 1, *n*_0_ = 0,*n*^*l*^(0) = 0 *g*_0_ = 1, *g*_*min*_ = 10^*−*6^, *g*_*max*_ = 70, 0.01 ≤ *α* ≤ 11, 0.2 ≤ *β* ≤ 10, Δ*α* = 0.2, and Δ*β* = 0.2.
- Figure 3. A) Numerical simulations of Model 2 with: *ϕ* = 50, *γ* = 2 *δ* = 4. The rest of the parameters are the same as in Fig. 2. B) Model 2, using equations equations 4 instead of equations 3 to simulate the dynamics of Abi sensors, with *γ* = 1 and *δ* = *−* 0.5. C) Numerical simulations of Model 2 with: *ϕ* = 50, *γ* = 5, and *δ* = 2 (light grey region) and with *ϕ* = 30, *γ* = 1.5, and *δ* = 0.5 (dark grey region). D) Same as in A.
- Figure 4. A) Same as Fig. 2. B-D) Numerical simulations of Model 3 for slow phages with *λ*_*c*_ = 1 and *µ*_*c*_ = 3. The rest of the parameters are the same as in Fig. 3.A.
- Figure 6. A) Same as in Fig. 2. B-D) Same as in Fig. 4.B.

